# Paired design of ATP bioluminescence method and colony counting method: whether it is suitable for evaluating the disinfection effect of environmental surface?

**DOI:** 10.1101/603860

**Authors:** Huiqiong Xu, Jiansheng Liang, Yimei Wang, Bin Wang, Tianbao Zhang, Xiaoli Liu, Lin Gong

## Abstract

ATP bioluminescence method has been used as a on-site rapid detection method in nosocomial infections control more and more. In this study, a paired design between four methods/detectors were performed to detect the environmental surface after disinfection. Two methods were ATP bioluminescence method and colony counting method (C), and ATP bioluminescence method included three kinds of detectors (B, P and N). Every surface was performed by two methods/detectors. ATP content per surface from ICU had no statistically significant difference with internal medicine wards of B and P, of which *p* was 0.435 and 0.260. By Spearman rank correlation coefficients, with the exception of better correlation between ATP content detected by B and P, the correlation between the remaining methods/detectors was poor or had no correlation. And the differences between detectors are statistically significant. Therefor, ATP bioluminescence method may not be suitable for the evaluation of disinfection effect, but be more appropriate for evaluating the cleanliness of environmental surface.

## Introduction

The environmental is one of the important links in the prevention and control of nosocomial infections. So, the cleaning quality and disinfection effect of it is of great significance to ensure the safety of patients and medical staff and reduce the risk of nosocomial infections [1-3]. Microbial culture (colony counting method) is the traditional and most common method to evaluate cleanliness and the disinfection effect, but it is time consuming. In the recent decade, most medical institutions, CDCs and health supervisions in China have begun to use ATP bioluminescence method to detect the surface disinfection of objects on-site widely. Because of the advantages of simplicity and rapidity, ATP bioluminescence method can solve the problem of hysteresis of colony counting method, and monitor the cleanliness of the ward environment according to the actual needs, so as to achieve “all staff, full-time, comprehensive” monitoring, to ensure that medical sites maintain a high standard of cleanliness for a long time, strengthen hospital cleanliness and reduce the chance of infection [4].

ATP bioluminescence method can detect the ATP content in the sample by biological fluorescence reaction using a luciferase assay and luminometer, in order to reflect the presence of microorganisms or other organic residues indirectly [5]. Since ATP bioluminescence technology was first described in 1963 by McElroy [6], the method has been introduced into food hygiene [7], nosocomial infections control and other fields. Our research team has used Staphylococcus aureus, Escherichia coli and Candida albicans as representative test bacteria of Gram-positive, Gram-negative and fungal, used the correlation coefficient (*r*) to describe the standard validity, and used the intra-class correlation coefficient (*ICC*) for reliability evaluation.. The results show that the ATP bio-fluorescence method has good accuracy and reliability. Some studies [8-10] show that ATP bioluminescence method has a good correlation with colony counting method. So, it has been widely used as a on-site rapid detection method for environmental sanitation disinfection, medical staff hand hygiene, medical device cleaning effect evaluation and so on in nosocomial infections control [11-14].

However, due to the absence of uniform evaluation standard, most medical institutions rely on the reference value provided by the manufacturer of testing detectors. In China, the relevant manufacturers recommend that the desktop ATP bioluminescence detector adopts RLU≤2 000 [15] as the qualified criterion for medical device cleaning, while the portable ATP bioluminescence detector standards vary, the smallest is RLU<30 [16], the largest is RLU≤500 [17]. Besides, ATP bioluminescence method has several limitations, especially the influencing factors. This makes the scientific and accurate judgment of the results to be questioned, which brings a certain difficulties to the majority of medical staffs in the daily related monitoring work. Therefore, our study aimed to compare ATP bioluminescence method with colony counting method, and also compare different kind of ATP bioluminescence detectors in monitoring the disinfection effect of environmental surface, to provide some data for answering “whether it is suitable for evaluating the disinfection effect of environmental surface?” and developing the unified judgment value.

## Methods

### Subjects

We performed environment surfaces from 22 medical institutions, including 12 tertiary hospitals.7 secondary hospitals and 3community health service centers. Of these hospitals, 13 are comprehensive hospitals, and the other 6 are specialized hospitals. Intensive care unit (ICU) and internal medicine wards were used as representatives of type II and type III environments, respectively. The samples mainly included treatment vehicles, treatment tables, bedside cabinets and doorknobs, etc. In total we performed 670 samples, 303 from ICU, and 367 from internal medicine wards. The study was carried out from 2017 through 2018.

### Disinfection and sampling Method

The environmental surfaces were disinfected by chlorine containing disinfectant of 500mg/L∼1000mg/L, then sampled after the surfaces became dry.

We sampled with swabs in ATP Surface Test or cotton swabs infiltrated by neutralizer (0.1% sodium thiosulfate). Surfaces evaluated in this study were collected by wiping a 100cm^2^ area with sterilized specification plate (5cm×5cm). If the total surface was smaller than 200cm^2^, each method or detector performed half of the area.

### ATP Bioluminescence Method

Three kinds of ATP bioluminescence detectors were selected which were used most in Wuhan, Hubei province. They were B (BT-112D, Beijing Chuangxin Shiji Biochemical Science&Technology Development Co.,Ltd., China) P (SystemSURE Plus, Hygiena, USA) and N (Clean-Trace NGi, 3M, USA). Two kinds of ATP surface test were used for the detector, with swab and some reagents. One was Clean-Trace™ ATP Surface Test (suitable for N, 3M, USA), and the other was UltraSnap™ ATP Surface Test (suitable for B and P, Hygiena, USA). We beat 20 times to make the swab and reagents mixed fully after sampling, then detected by the detector to get the RLU value. The results were reported as ATP content (mol) per surface, which was converted by standard curve.

First, the ATP standard solution (100nmol, BioThema, Sweden) was diluted to graded concentration by pure water, which were 1.0×10^−7^mol/L, 5.0×10^−8^mol/L, 1.0×10^−8^mol/L, 5.0×10^−9^mol/L, 1.0×10^−9^mol/L, 5.0×10^−10^mol/L, 1.0×10^−10^mol/L, 5.0×10^−11^mol/L, 1.0×10^−11^mol/L, 5.0×10^−12^mol/L and 1.0×10^−12^mol/L. Then, each concentration was taken 10μl to drop onto the swab and RLU value was gotten following the step above. We repeated 3 times for each concentration and took the average. Finally, the standard curve of x and y axes were fitted with the base 10 logarithms of ATP content (10^−17^mol) and RLU, which was showed y=ax+b. Each detector had its own standard curve.

### Colony Counting Method

Colony counting method was a classical microbial method. After sampling, The nutritional agar culture medium (Qingdao Hope Bio-technology CO.,LTD., China) was used to monitor microbial contamination on each surface. The experimental process was according to Hygienic standard for disinfection in hospitals (GB15982-2012) [18]. The total colonies was estimated as CFU per surface, which equaled to the average colonies per dish multiplied by the dilution multiples of the sample solution.

### Study Design

There were two methods, ATP bioluminescence method and colony counting method (C). Besides, ATP bioluminescence method included three kinds of detectors (B, P and N). So, in this study, the paired design was designed between these four methods/detectors. Every surface was performed by two methods/detectors. Every time, the two methods/detectors were used to sample different points of the same surface.

### Statistical Analysis

All of the detecting data were transferred to Microsoft Office Excel, and statistical analysis were conducted using SPSS version 16.0, with a p value of <0.05 used to be considered as statistically significant. All the results were given separately according to different resource (ICU and internal medicine wards). And analysis was done using the nonparametric test. Rank correlation was used to compare CFU and ATP content or ATP contents by different detectors, and Wilcoxon singned-rank test was used to compare ATP contents between different detectors.

## Results

### Standard curve

In this study, we used 3 B, 3 P and 6 N. Of the standard curve experiment, the desktop detector (B) could detect to 1.0×10^−17^mol, and the portable detector (P and N) could detect to 1.0×10^−15^mol. The standard curve and linear correlation coefficient (*r)* of each detectors are shown in Table 1.

**Table 1.** The Standard Curve and Linear Correlation Coefficient (*r)* of Each Detectors.

| Detector | Standard curve<sup>a</sup> | $r$ | Detector | Standard curve<sup>a</sup> | $r$ |
| --- | --- | --- | --- | --- | --- |
| B (1) | $y=0.893x+0.990$ | 0.987 | N (1) | $y=0.914x-0.785$ | 0.996 |
| B (2) | $y=0.987x+1.378$ | 0.999 | N (2) | $y=0.905x-0.761$ | 0.996 |
| B (3) | $y=1.004x+0.862$ | 0.998 | N (3) | $y=0.931x-0.852$ | 0.998 |
| H (1) | $y=0.941x-1.714$ | 0.990 | N (4) | $y=0.874x-0.409$ | 0.995 |
| H (2) | $y=1.022x-0.241$ | 0.996 | N (5) | $y=0.851x-0.356$ | 0.994 |
| H (3) | $y=1.002x-1.992$ | 0.998 | N (6) | $y=0.853x-0.388$ | 0.994 |

<sup>a</sup> x and y are fitted with the base 10 logarithms of ATP content ( $10^{-17}$ mol) and RLU.

### Distribution of CFU/ATP content

By Kolomogorov-Smirnov test, the distribution of CFU and ATP content in this study were not obey the normal distribution, with a *p* value of <0.05. So, we used median and inter-quartile range (*Q*) to describe the central tendency and discrete tendency, respectively (Table 2). In this study, 286 cases of C (88.0%) were less than or equal to 10 CFU/surface, 218 (67.1%) cases of C were 0 CFU/surface. According to GB15982-2012 [18], 276 cases of C (84.9%) met the hygienic standard, which required ICU with a CFU value of ≤5 and internal medicine wards with a CFU value of ≤10. By Wilcoxon rank sum test, CFU/ATP content per surface from ICU was lower than internal medicine wards of C and N, of which *p* was 0.024 and 0.002, respectively. However, ATP content per surface from ICU had no statistically significant difference with internal medicine wards of B and P, of which *p* was 0.435 and 0.260.

**Table 2.**
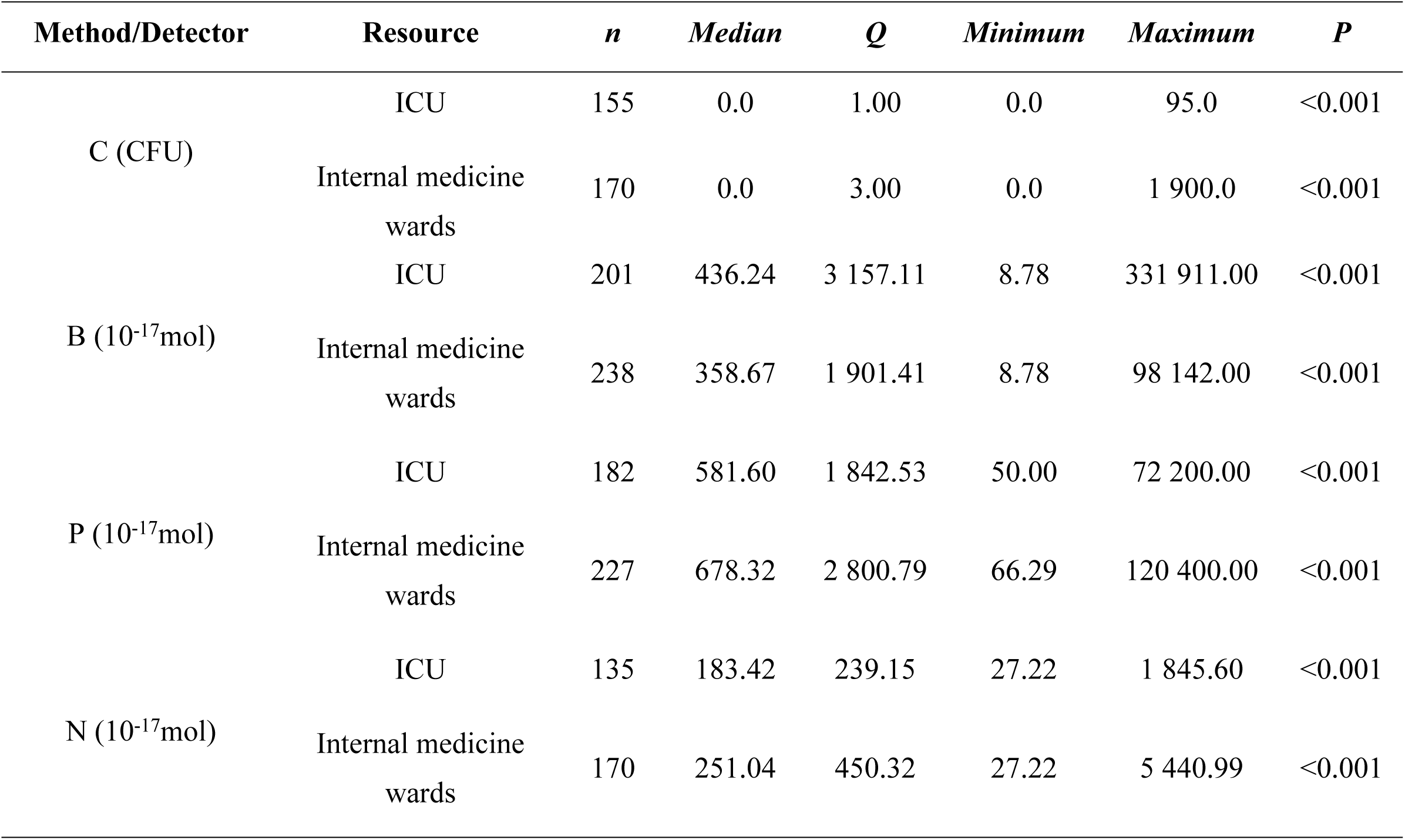
The Distribution of CFU/ATP Content (per Surface) after Disinfection.

| Method/Detector | Resource | *n* | *Median* | *Q* | *Minimum* | *Maximum* | *P* |
| --- | --- | --- | --- | --- | --- | --- | --- |
| C (CFU) | ICU | 155 | 0.0 | 1.00 | 0.0 | 95.0 | $<0.001$ |
| | Internal medicine wards | 170 | 0.0 | 3.00 | 0.0 | 1 900.0 | $<0.001$ |
| B ($10^{-17}$mol) | ICU | 201 | 436.24 | 3 157.11 | 8.78 | 331 911.00 | $<0.001$ |
| | Internal medicine wards | 238 | 358.67 | 1 901.41 | 8.78 | 98 142.00 | $<0.001$ |
| P ($10^{-17}$mol) | ICU | 182 | 581.60 | 1 842.53 | 50.00 | 72 200.00 | $<0.001$ |
| | Internal medicine wards | 227 | 678.32 | 2 800.79 | 66.29 | 120 400.00 | $<0.001$ |
| N ($10^{-17}$mol) | ICU | 135 | 183.42 | 239.15 | 27.22 | 1 845.60 | $<0.001$ |
| | Internal medicine wards | 170 | 251.04 | 450.32 | 27.22 | 5 440.99 | $<0.001$ |

### Correlation of different methods/detectors

By Spearman rank correlation coefficients (*r*_*s*_), with the exception of better correlation between ATP content detected by B and P, the correlation between the remaining methods/detectors was poor or had no correlation (Table 3).

**Table 3.** The Correlation of CFU/ATP Content (per Surface) between Methods/Detectors.

| Method/<br>Detector | C (CFU) |  |  | B (10<sup>-17</sup>mol) |  |  | P (10<sup>-17</sup>mol) |  |  | N (10<sup>-17</sup>mol) |  |  |
| --- | --- | --- | --- | --- | --- | --- | --- | --- | --- | --- | --- | --- |
| | $r_s$ | $P$ | $n$ | $r_s$ | $P$ | $n$ | $r_s$ | $P$ | $n$ | $r_s$ | $P$ | $n$ |
| C (CFU) | - | - | - | 0.293<sup>a</sup> | 0.005<sup>a</sup> | 90<sup>a</sup> | 0.244<sup>a</sup> | 0.029<sup>a</sup> | 80<sup>a</sup> | 0.313<sup>a</sup> | 0.049<sup>a</sup> | 40<sup>a</sup> |
| B (10<sup>-17</sup>mol) | 0.450<sup>b</sup> | <0.001<sup>b</sup> | 93<sup>b</sup> | - | - | - | 0.904<sup>a</sup> | <0.001<sup>a</sup> | 121<sup>a</sup> | 0.246<sup>a</sup> | 0.062<sup>a</sup> | 58<sup>a</sup> |
| H (10<sup>-17</sup>mol) | 0.485<sup>b</sup> | <0.001<sup>b</sup> | 90<sup>b</sup> | 0.931<sup>b</sup> | <0.001<sup>b</sup> | 139<sup>b</sup> | - | - | - | 0.073<sup>a</sup> | 0.613<sup>a</sup> | 51<sup>a</sup> |
| N (10<sup>-17</sup>mol) | 0.564<sup>b</sup> | <0.001<sup>b</sup> | 40<sup>b</sup> | 0.378<sup>b</sup> | <0.001<sup>b</sup> | 81<sup>b</sup> | 0.466<sup>b</sup> | <0.001<sup>b</sup> | 70<sup>b</sup> | - | - | - |

<sup>a</sup> It represents the result of ICU.
<sup>b</sup> It represents the result of internal medicine wards.

### Compare between detectors

By Wilcoxon singned-rank test, the ATP content of ICU environment surface detected by P and N had no statistically significant difference, while all of the other ATP content detected by different detectors showed a statistically significant difference (Table 4).

**Table 4.**
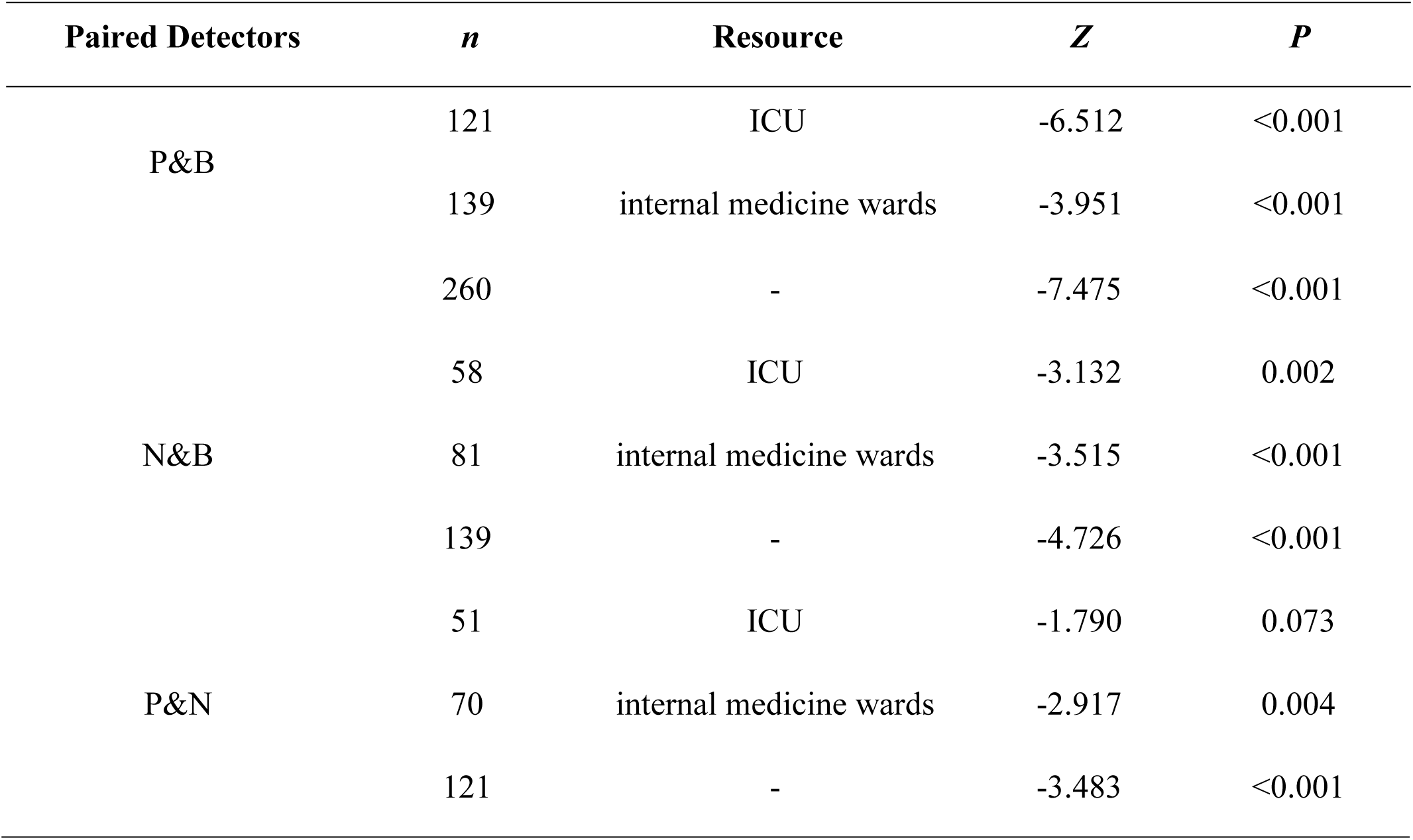
The Comparison of ATP Content (10^−17^mol) Detected by Different Detectors.

## Discussion

Contamination of hospital surfaces plays an important role in the transmission and diffusion of several pathogens, which may infect patients, then contaminate the hands of medical staffs, and be passed on to other patients, or contaminate other surfaces. This circulation may lead to the occurrence or even outbreak of nosocomial infections. Consequently, methods to assess hospital environments cleaning can be considered an integral part of infections prevention and control programs [19]. At present, the most used methods are visual inspection, microbial culture, fluorescent markers and ATP bioluminescence method, which are the main tools and methods identified by CDC of USA [20]. Nowadays, ATP bioluminescence method is increasingly used to evaluate the effect of environmental surfaces cleaning and disinfection. An independent study [21] evaluating the potential role of the ATP tool in assessing cleaning practice, while noting several limitations in the ATP system, concluded that the tool could potentially be used effectively for environment service education. Boyce JM [22, 23] reported monitoring the effectiveness by ATP bioluminescence assay could significant improve the daily cleaning.

The RLU value of different detectors always varies with difference in photometric capability between photomultiplier tubes, difference between reagent sensitivity, and so on. But in theory, the ATP content of microorganism is certain, so the difference between different detectors should not be statistically significant after converting the RLU value into ATP content (mol). We try to use standard curve to convert the RLU value into ATP content, and then compare the ATP content between different detectors. The results show that, after disinfection, the correlation between the ATP content and CFU is poor, and the correlation between the ATP content of three detectors is also not ideal(table 3), which may be related to the low detection rate of microbial culture. In our study, 88% of the colony counting method are less than or equal to 10 CFU/surface, even 67% are 0 CFU/surface. In 2003, Hattori et al. detected 54 kinds of microorganisms, the ATP contents of the gram-negative bacteria, gram-positive bacteria and yeasts ranged from 0.40 to 2.70×10^−18^mol/CFU (mean=1.5×10^−18^mol/CFU), from 0.41 to 16.7×10^−18^mol/CFU (mean=5.5×10^−18^mol/CFU), and from 0.714 to 54.6×10^−18^mol/CFU (mean=8.00×10^−18^mol/CFU), respectively [24]. According to this, more than four-fifth of the ATP content of our study are around 10^−17^mol, but the detectors used in our study could only detect above 10^−17^mol (B) and 10^−15^mol (P and N). So, the surface after disinfection that is too clean may not be suitable for evaluating by ATP bioluminescence method. Besides, the differences between different detectors are basically statistically significant (Table 4). Maybe it has something to do with the homogeneity of the samples. And fluctuating could also be caused by the presence of chemicals and other materials, such as the residues of detergent or disinfectants, microfiber products, and manufactured plastics [19, 25]. The ATP detection value may also vary depending on the raw material composition of the detected object [26].

There are a number of limitations to our study. ATP bioluminescence method can detect all organic material, including bacteria, blood, excretions, human secretions, food and so on [27]. So, in order to evaluate the disinfection effect more accurate, we need to do a good job of cleaning before disinfection. But in our study, we haven’t specifically emphasized the role of cleaning. However, on the other hand, it is precisely because ATP bioluminescence method can not only detect the ATP of microorganisms, but also detect organic matters, so it may not be suitable for the evaluation of disinfection effects, but be more appropriate for evaluating the cleanliness, or as an early warning method of microbial contamination. Next, we will continue to study the assessment of the surface cleanliness of environmental objects to provide more data support for the criteria of the ATP bioluminescence method.

## Acknowledgments

We are grateful to all authors of the literature included in this article for supporting material and helpful information. We also would like to thank Hubei Center for Disease Control and Prevention (Xiaobo Huang), Wuhan No. 1 Hospital (Zhigang Liu, Yanqiong Peng), Wuhan Third Hospital (Xiaoting Chen, Puqin Tang), the Central Hospital of Wuhan (Xiaoman He, Juhong Qiu), Hubei Maternal and Child Health Hospital (Xinyun Lei, Xia Gao), the Eighth Hospital of Wuhan (Jianrong Tang, Qiuming Zhu), Beijing Chuangxin Shiji Biochemical Science&Technology Development Co., Ltd.(Zhi Wang), and 3M China (Jifeng Zhang) for providing assistance with data acquisition.

